## Supplemental Figures for "Autoinhibited kinesin-1 adopts a hierarchical folding pattern"

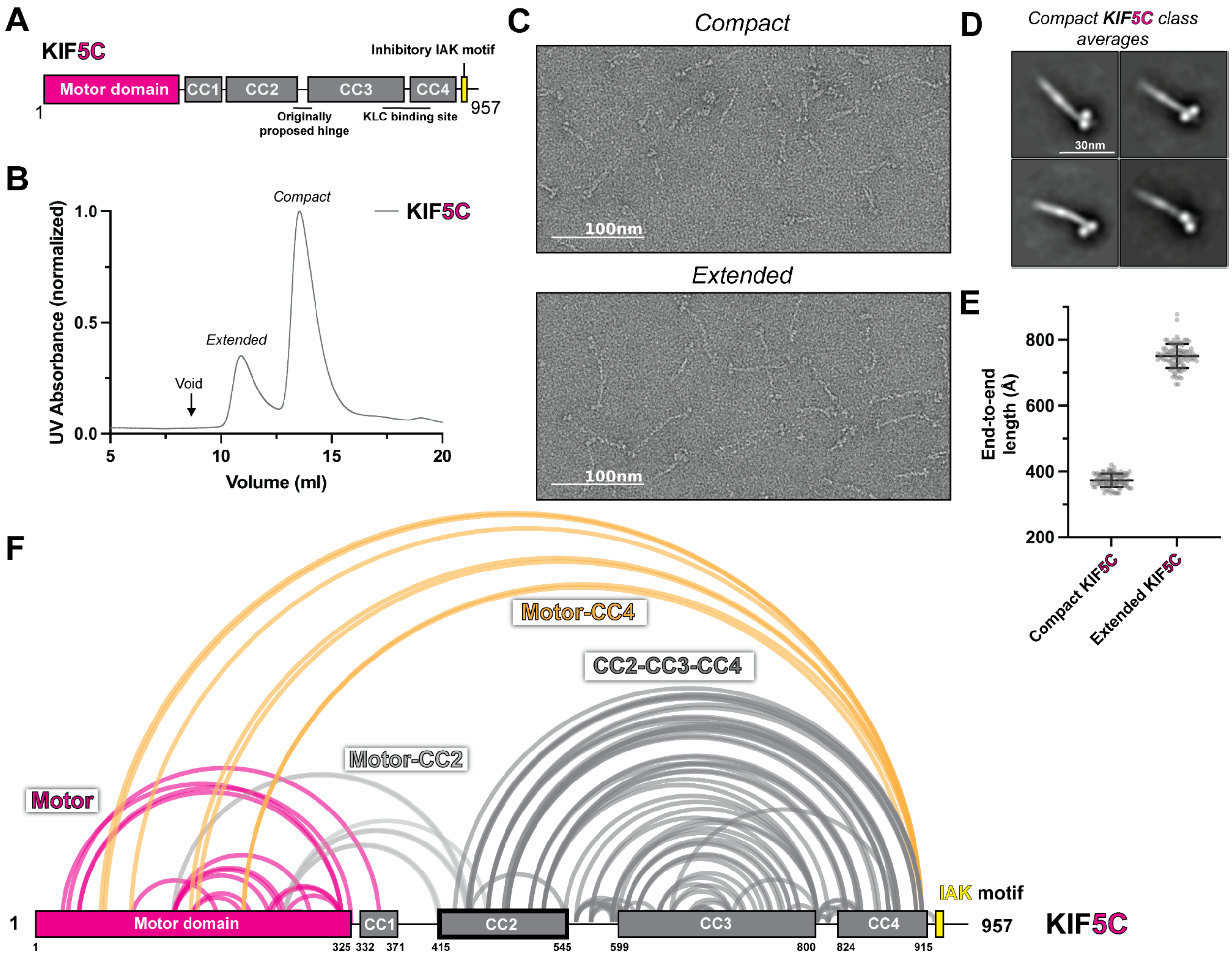


**Figure S1. KIF5C adopts a hierarchical folding pattern.**

(A) The domain diagram of KIF5C. (B) The size exclusion chromatography profile of KIF5C. (C) The example micrographs from KIF5C negative staining EM. (D) Class averages of compact KIF5C. (E) End to end length measurements of KIF5C in two states (N=100). (F) Crosslinked lysine pairs in KIF5C were mapped onto the domain diagram and divided into four groups.

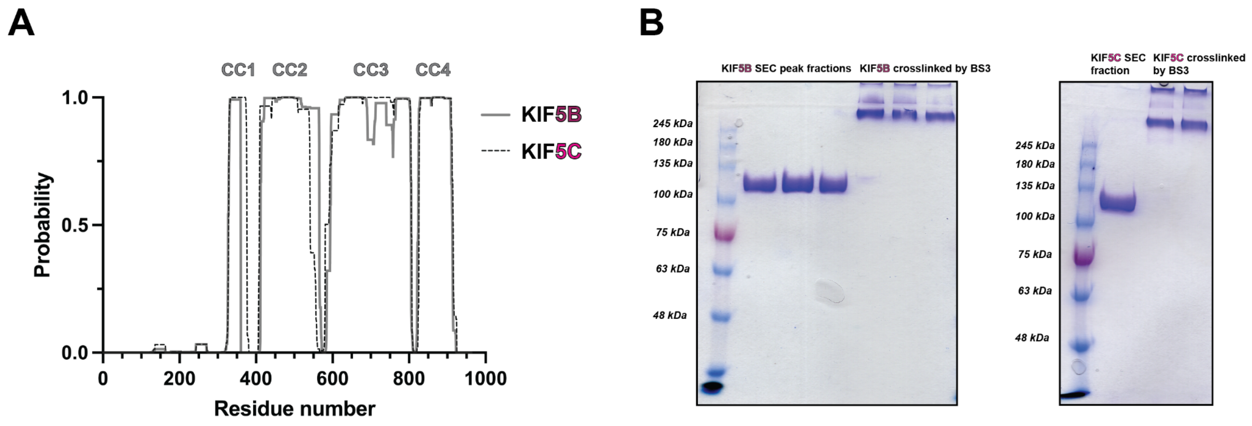


**Figure S2. Structural analysis and purification of KIF5B and KIF5C.**
(A) Coiled-coil prediction results from Marcoil (Delorenzi & Speed, 2002) on MPI Bioinformatics Toolkit (Gabler et al., 2020; Zimmermann et al., 2018). (B) SDS-PAGE analysis of crosslinked KIF5B and KIF5C samples.

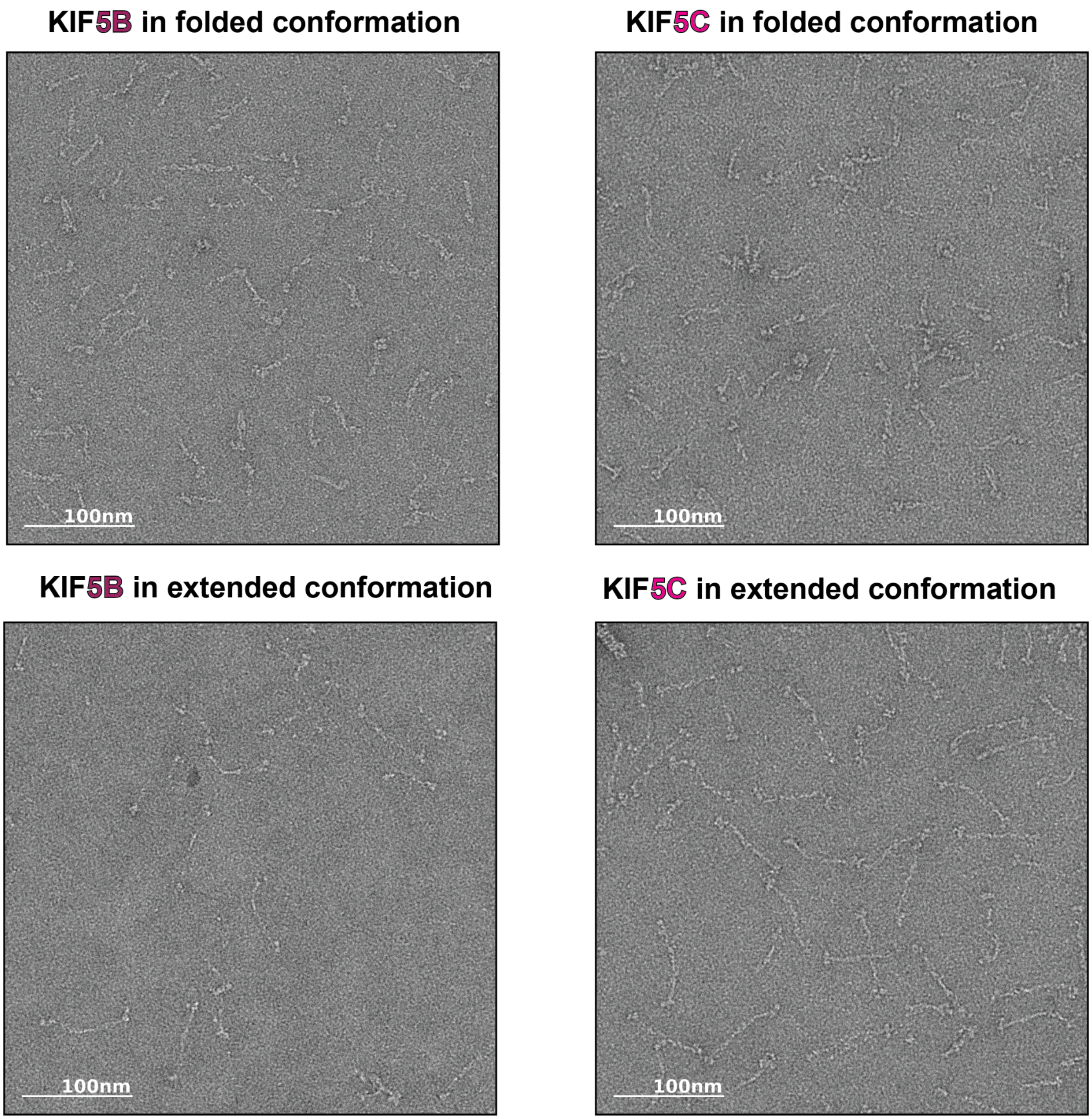


**Figure S3. Negative stain EM of KIF5B and KIF5C.**
Representative micrographs of KIF5B and KIF5C in extended and compact states.

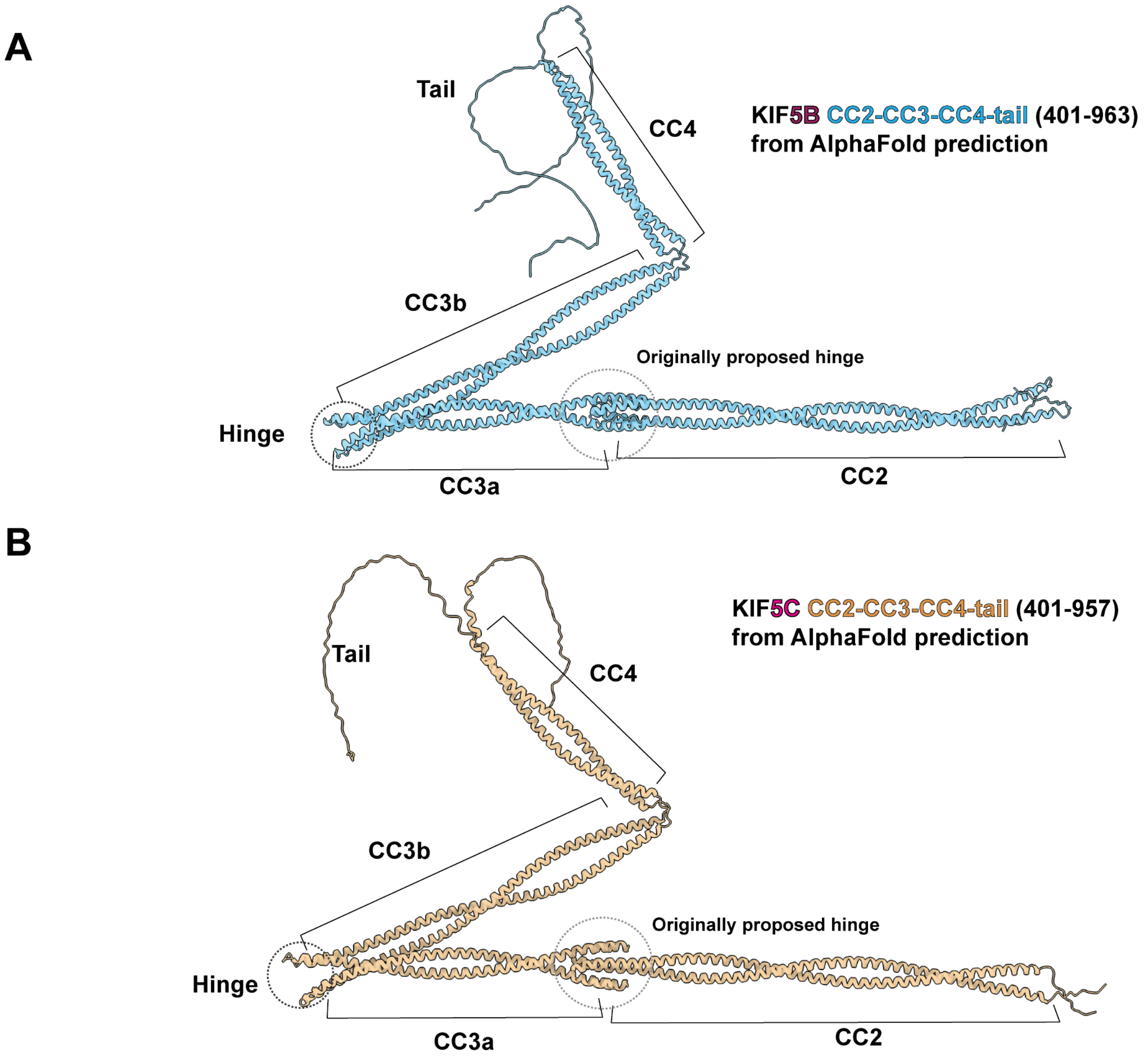


**Figure S4. The AlphaFold predicted structures of KIF5B and KIF5C stalk and tail.**

(A) The predicted structure of KIF5B (401-963). (B) The predicted structure of KIF5C (401-957). The coiled-coil prediction results and location of the originally proposed hinge were mapped onto the structures. The newly identified hinge sits in between the CC3a and CC3b.

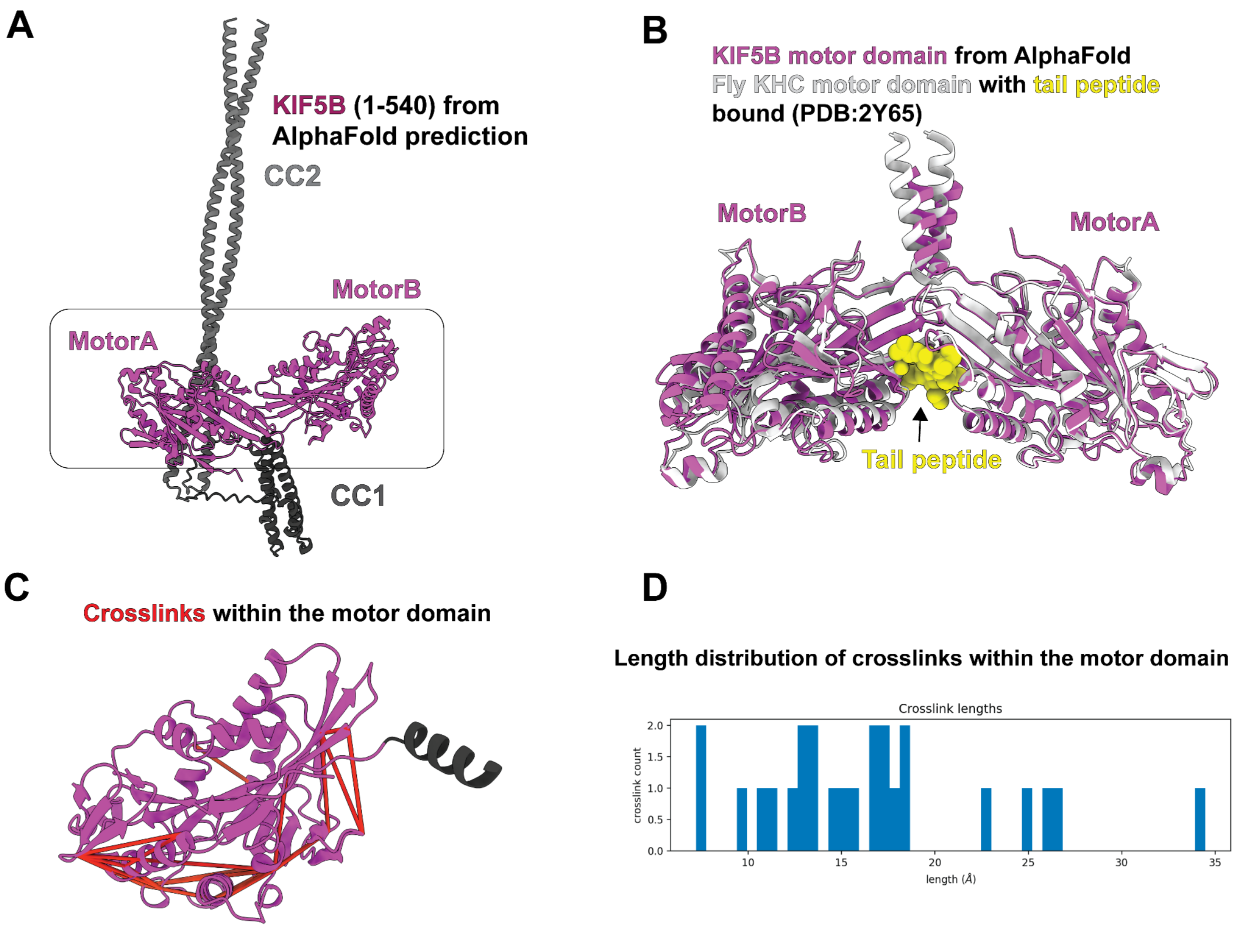


#### Figure S5. Predicted KIF5B motor domain resembles the tail peptide bound state.

(A) The predicted structure of KIF5B (1-540) containing motor domain, CC1 and CC2. (B) Superposition of predicted KIF5B motor domain structure (purple) with tail peptide bound fly KHC motor domain structure (light gray) (PDB:2Y65). (C) Intra-motor domain crosslinks were mapped to the predicted motor domain structure. (D) The length distribution of crosslinks within the motor domain.

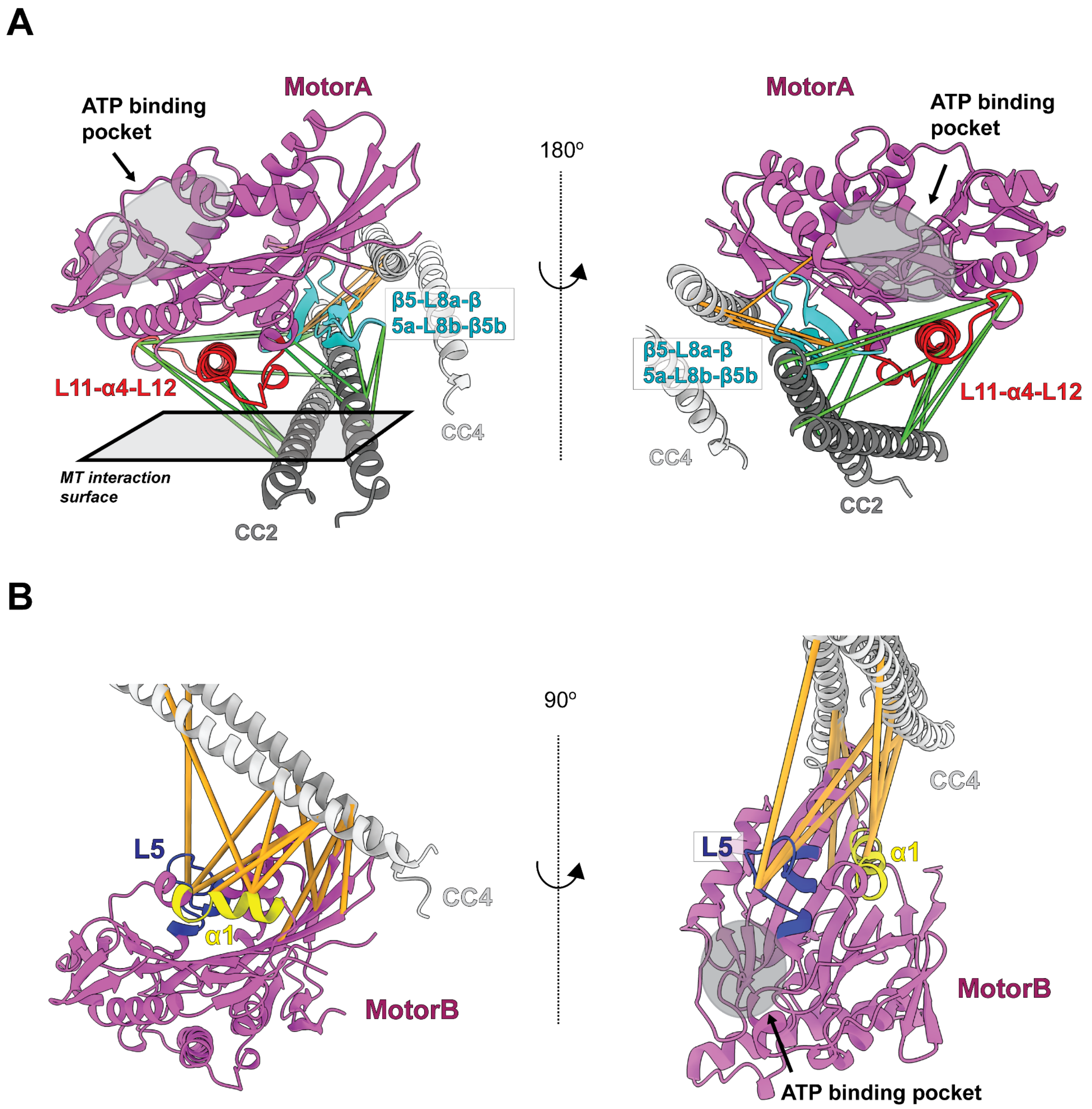


**Figure S6. The stalk crosslinks to multiple points on the motor domain.**
(A) The microtubules binding interface in motorA was crosslinked to the CC2 and CC4. The red colored region is L11-α4-L12, mainly for interacting with α-tubulin. The Cyan colored region is β5-L8, which is responsible for interacting with β-tubulin. (B) Two elements (L5 and α1) in motorB were crosslinked to CC4.

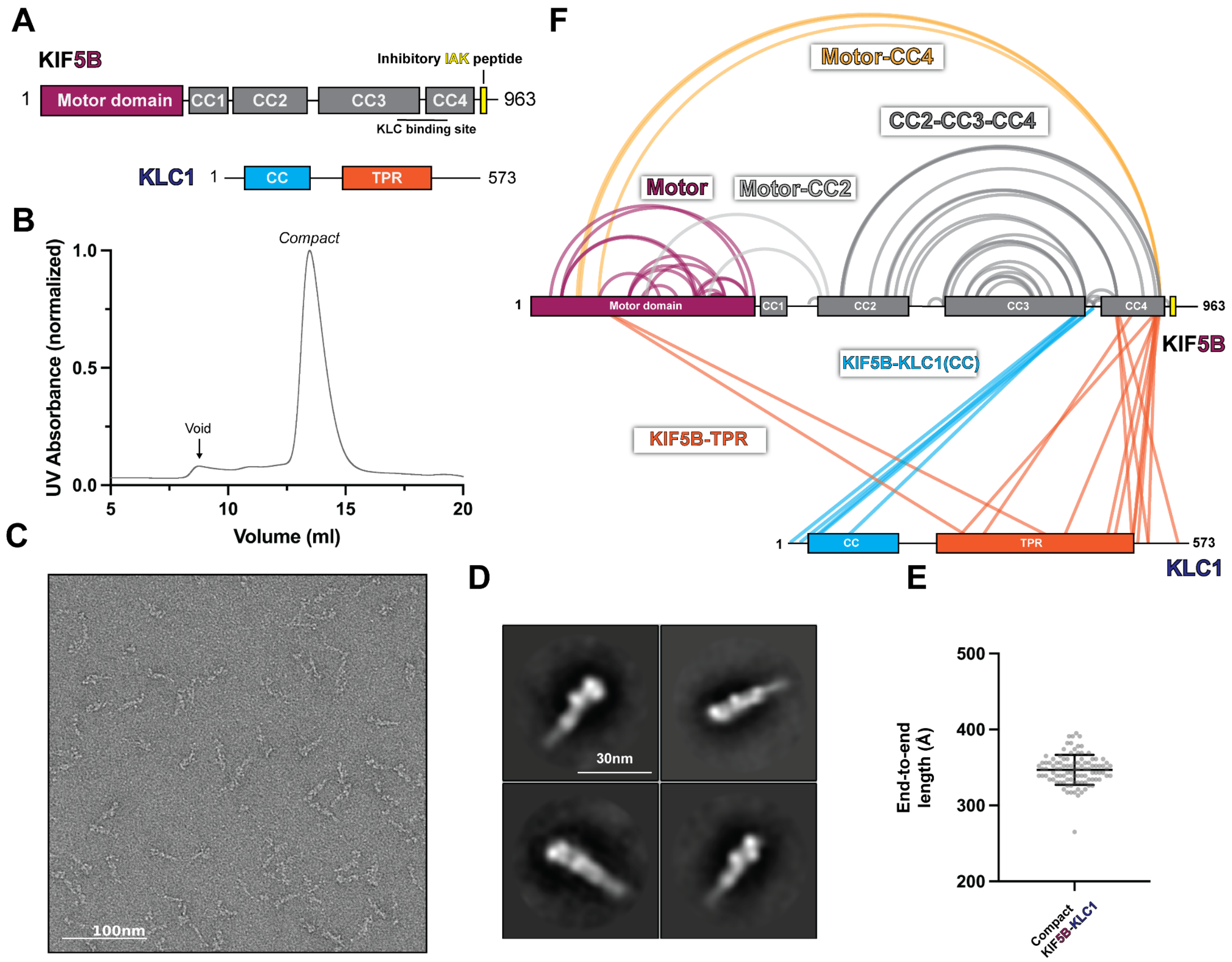


### Figure S7. KLC1 maintains the folding pattern of KIF5B.

(A) The domain diagram of KIF5B-KLC1. (B) The size exclusion chromatography profile of KIF5B-KLC1. (C) The example negative staining EM micrograph of KIF5B-KLC1 in compact state. (D) Examples of 2D class averages of KIF5B-KLC1. (E) End to end length measurements of KIF5B-KLC1 in compact state (N=100). (F) Crosslinked lysine pairs in KIF5B-KLC1, shown in the domain diagram.

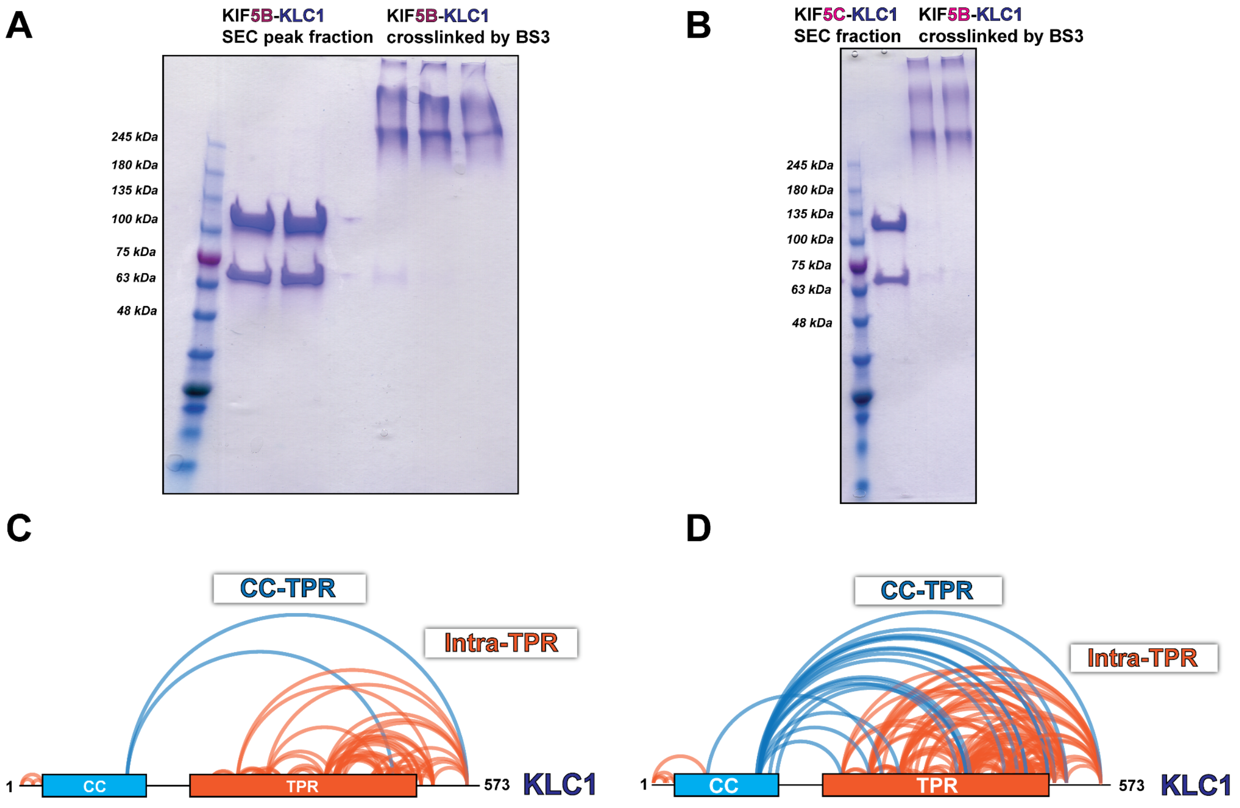


**Figure S8. SDS-PAGE analysis of kinesin-1 heterotetramer (KIF5B-KLC1 and KIF5C-KLC1) and the intra-KLC1 crosslinks.**

(A) SDS-PAGE analysis of KIF5B-KLC1. (B) SDS-PAGE analysis of KIF5C-KLC1. (C) Intra-KLC1 crosslinks from KIF5B-KLC1. (D) Intra-KLC1 crosslinks from KIF5C-KLC1.

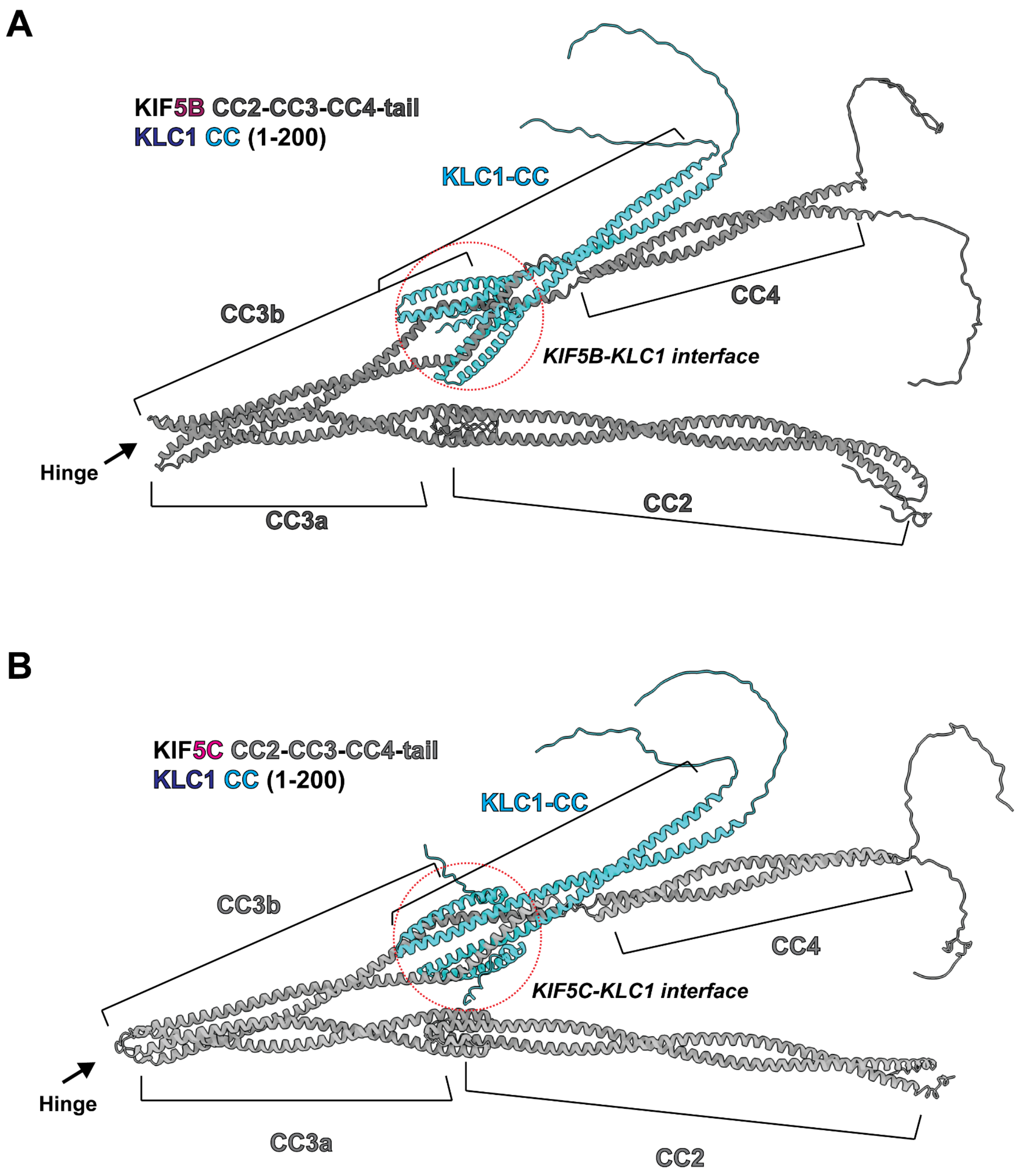


**Figure S9. The AlphaFold predicted structure of KIF5B/C stalk and KLC1.**
(A) The predicted structure of KIF5B (401-963) - KLC1 (1-200). (B) The predicted structure of KIF5C (401-957) - KLC1 (1-200). The coiled-coil prediction results and location of binding site between kinesin heavy chain and light chain were mapped onto the structure.

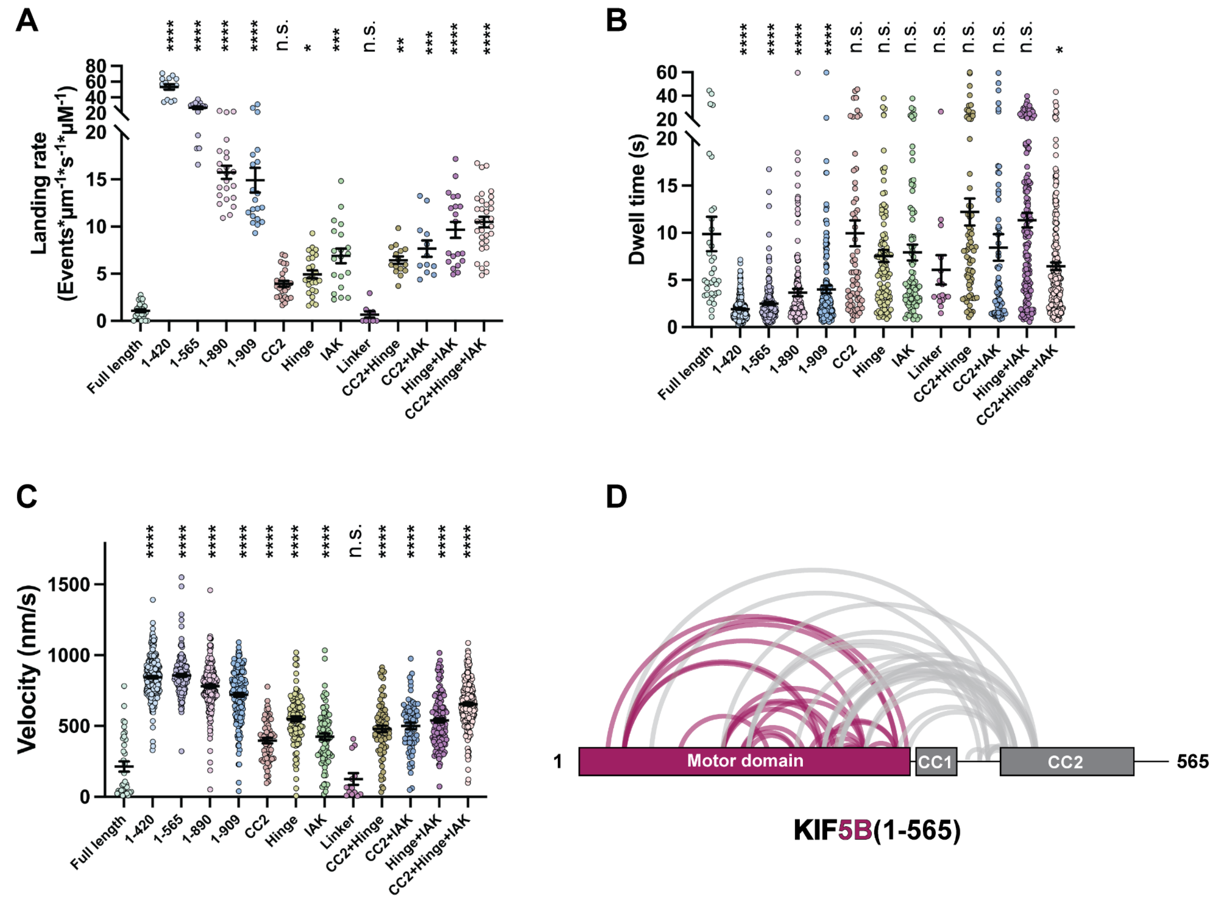


#### Figure S10. The landing rate, dwell time and velocity distribution of all KIF5B variants and the XL-MS results of KIF5B (1-565).

(A) The landing rate distribution of KIF5B with different mutations and truncations. Landing rate (Events/μm/s/μM): Lines show mean ± SEM: 1.07 ± 0.16 (full length), 53.42 ± 3.47 (1-420), 26.87 ± 1.65 (1-565), 15.75 ± 0.70 (1-890), 14.92 ± 1.30 (1-909), 3.93 ± 0.32 (CC2), 4.93 ± 0.42 (Hinge), 6.90 ± 0.78 (IAK), 0.67 ± 0.32 (Linker), 6.44 ± 0.40 (CC2+hinge), 7.66 ± 0.86 (CC2+IAK), 9.65 ± 0.85 (Hinge+IAK), 10.49 ± 0.57 (CC2+hinge+IAK). N=2, n=25, 14, 15, 23, 20, 25, 25, 20, 9, 15, 12, 20, and 32 MTs. One-way ANOVA followed by Dunnett's test. *P<0.05, **P<0.01, ***P<0.001, ****P<0.0001, n.s not significant. (B) The dwell time distribution of KIF5B with different mutations and truncations. Dwell time (s): Lines show mean ± SEM: 9.88 ± 1.83 (full length), 1.91 ± 0.08 (1-420), 2.49 ± 0.16 (1-565), 3.67 ± 0.40 (1-890), 3.99 ± 0.41 (1-909), 9.93 ± 1.37 (CC2), 7.54 ± 0.64 (Hinge), 7.91 ± 0.85 (IAK), 6.07 ± 1.55 (Linker), 12.21 ± 1.42 (CC2+hinge), 8.45 ± 1.42 (CC2+IAK), 11.34 ± 0.76 (Hinge+IAK), 6.46 ± 0.41 (CC2+hinge+IAK). N=2, n= 36, 253, 185, 182, 180, 61, 103, 83, 16, 75, 68, 143, and 223 events. One-way ANOVA followed by Dunnett's test. *P<0.05, ****P<0.0001, n.s not significant. (C) The velocity distribution of KIF5B with different mutations and truncations. Velocity (nm/s): Line shown mean ± SEM: 241.2 ± 36.04 (full length), 844.5 ± 8.89 (1-420), 858.0 ± 9.72 (1-565), 781.3 ± 12.47 (1-890), 720.9 ± 13.75 (1-909), 396.7 ± 19.15 (CC2), 549.7 ± 17.94 (Hinge), 424.1 ± 21.86 (IAK), 125.8 ± 40.93 (Linker), 479.4 ± 22.83 (CC2+Hinge), 500.1 ± 22.4 (CC2+IAK), 539.4 ± 15.24 (Hinge+IAK), 654.7 ± 10.94 (CC2+Hinge+IAK). N=2, n= 36, 253, 185, 182, 180, 61, 103, 83, 13, 75, 68, 143 and 223 events. One-way ANOVA followed by Dunnett's test. ****P<0.0001, n.s not significant. (D) The crosslinked pairs in KIF5B (1-565), suggesting that motor-CC2 folding is present in truncated kinesin and is independent of stalk folding.

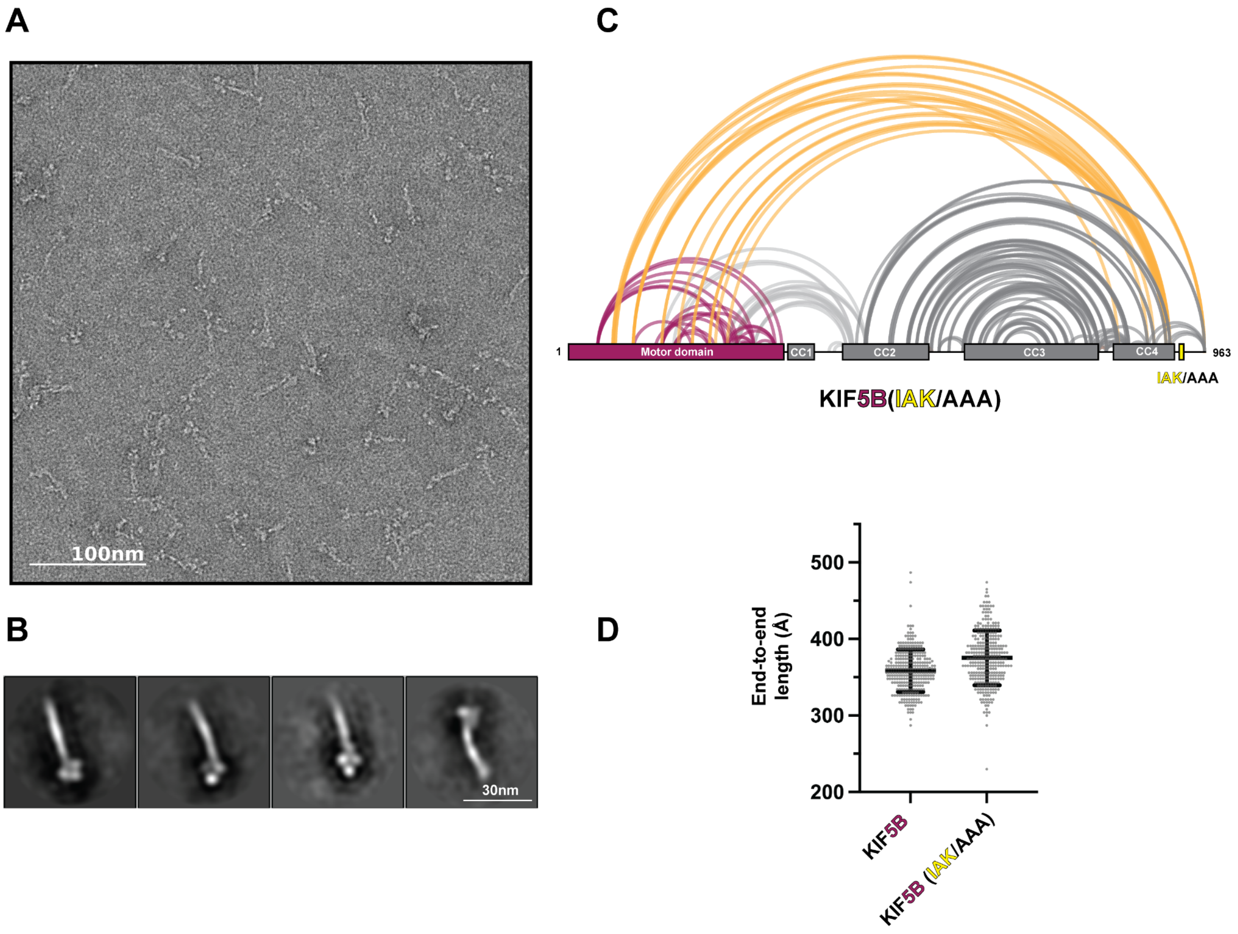


**Figure S11. IAK mutation does not relieve KIF5B folding.**
(A) Negative staining EM of crosslinked KIF5B(IAK/AAA). (B) The example class averages of KIF5B(IAK/AAA). (C) The crosslinked pairs in KIF5B(IAK/AAA) from XL-MS. (D) End-to-end length measurement of KIF5B(WT) and KIF5B(IAK/AAA). (N=300)

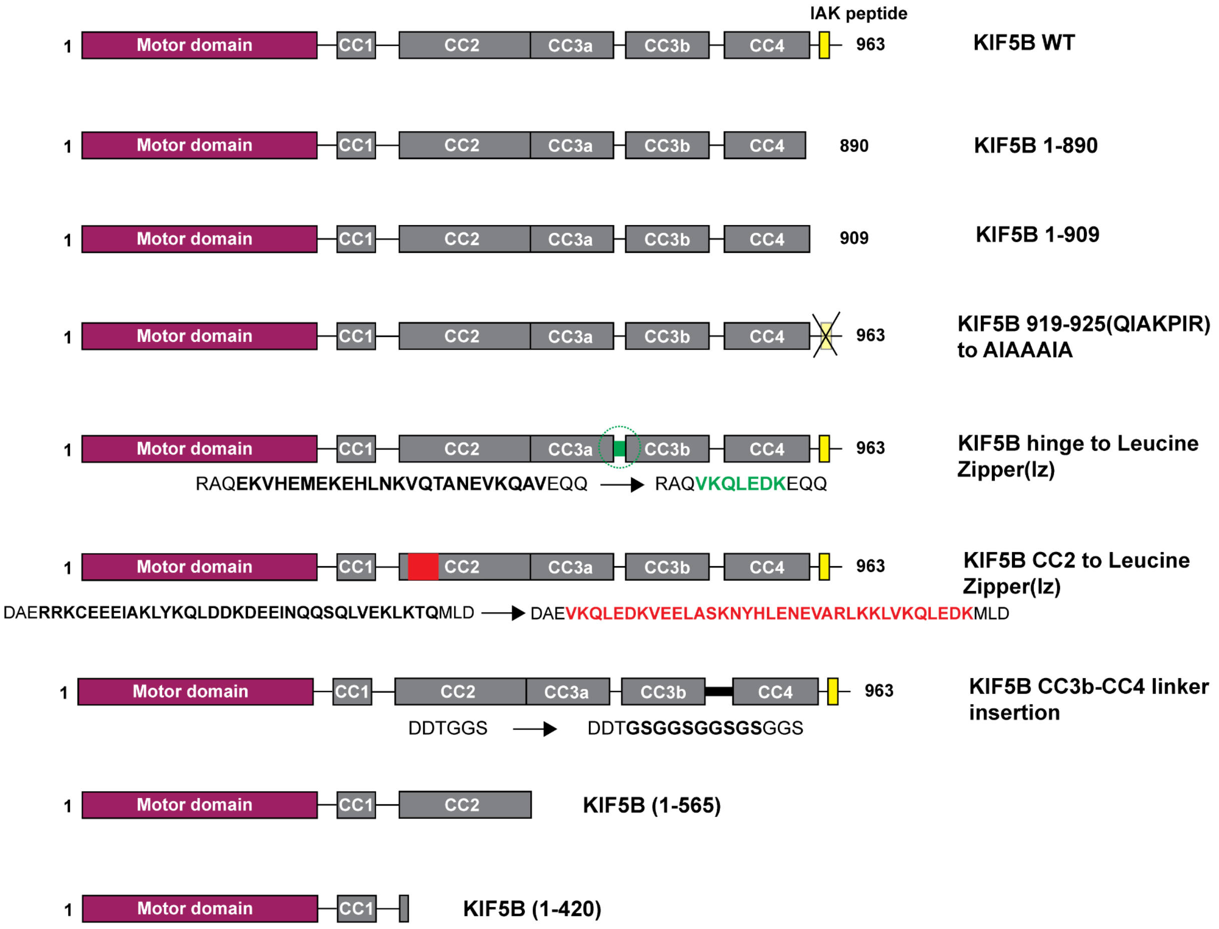


**Figure S12. Description of KIF5B constructs used in the single molecule motility assay.**
